# Investigation of co-encapsulation of pancreatic beta cells and curcumin within alginate microcapsules

**DOI:** 10.1101/2023.02.25.530032

**Authors:** Zahra Hosseinzadeh, Iran Alemzadeh, Manouchehr Vossoughi

## Abstract

Cell encapsulation is an ideal approach for the replacement of pancreatic function in Type 1 diabetes. Poor biocompatibility of microcapsules generates an inflammatory response in the implantation site and induces fibrosis infiltration, which causes microencapsulated cell death and graft failure. To prevent inflammation after implantation, composite microcapsules which exhibit anti-inflammatory properties were designed. This study is about co-encapsulating beta cells and curcumin within 1.5% alginate by the jet-breaking regime of the syringe pump. The microcapsules size distribution and rate of the alginate solution were characterized to find uniform particles. Micro-size particles were attained at a rate of 25 ml/min. Uniform spherical microcapsules (200–300 μm) were created in large amounts in a short period. Microcapsule breakage was less than 3% during 7 days and demonstrated the stability of the encapsulation method. Insulin secretion and cell viability assays were performed 1, 3, and 7 days after microencapsulation by GSIS and MTT assays. No significant differences in the amount of insulin secretion and beta cell viability were observed among free cells, alginate microcapsules, and curcumin-alginate microcapsules during 7 days (P > 0.05). Therefore, curcumin and alginate membrane did not show any harmful impacts on the function and survival of the beta cells.

## 2. INTRODUCTION

Type 1 diabetes mellitus (T1DM) is an autoimmune and degenerative disease that is caused by the destruction of insulin□producing pancreatic β cells due to complex genetic and environmental factors interactions (Atkinson, Campbell-Thompson, Kusmartseva, & Kaestner, 2020; Holt, Lupsa, Lee, Bassyouni, & Peery, 2022; Jara et al., 2020; Schaschkow et al., 2020). Diabetes patients suffer from insulin deficiency and high blood glucose levels (hyperglycemia). Daily insulin injection is the first therapy for T1DM that does not mimic the beta cells’ insulin release patterns. Thus, the patients may encounter hyperglycemia or hypoglycemia, which causes cognitive impairment, seizures, retinopathy, kidney diseases, and coma (Espona-Noguera et al., 2019; Li et al., 2014; Nikravesh, Cox, Ellis, & Grover, 2017; Qi, 2014; Skrzypek et al., 2017). The other therapy is to transplant the whole pancreatic system. However, the problems with this approach are the scarcity of pancreas donors and using a strong immunosuppressive regimen (Espona-Noguera et al., 2019).

The limitations above mentioned have led to the exploration of alternative approaches for islet transplantation. One approach to solving these problems is providing a 3D environment and encapsulating beta cells within microcapsules (Schaschkow et al., 2020; Toda et al., 2019). The microcapsule allows small molecules like oxygen, glucose, and insulin penetration. However, it does not allow larger components, such as natural killer cells, dendritic cells, and macrophages, to penetrate the capsule and destroy the cells (Bansal, D’Sa, & D’Souza, 2019; Ghoneim, Refaie, Elbassiouny, Gabr, & Zakaria, 2020; Kim et al., 2021). This technology encapsulates beta cells within a biocompatible microcapsule, which provides an immuno-isolated environment. Therefore, when the beta cells were implanted in the body, this immuno-isolation space can decline the use of immunosuppressive drugs. In addition, this strategy solves the lack of pancreas donors by employing the transplantation of heterologous islets and insulin-secreting cells (Christoffersson, 2022; Mochizuki et al., 2020). Encapsulation of cells within biomaterials has the potential to enhance immunosuppression. While cell encapsulating has been an examined strategy to prevent cell-cell interaction, antigen recognition, and immune cell attack, in vivo results represented that it is inadequate to block indirect immune system activities (Samojlik & Stabler, 2021).

Many materials such as alginate, collagen, agarose, cellulose, chitosan, gelatin, PEG, and silica have been mainly used for microencapsulation, while most of them show weak performance in beta cell encapsulation (Espona-Noguera et al., 2019; Toda et al., 2019). Among them, alginate is commonly used for cell microencapsulation because it exhibits high biocompatibility for encapsulated cells (Chan, Krishnan, Alexander, & Lakey, 2017; de Vos, Faas, Strand, & Calafiore, 2006). Alginate does not reveal toxic effects on encapsulated cells; however, its insufficient biocompatibility causes fibrosis on the microcapsule’s surface in the implantation site of the body, which is generated by the inflammatory response via macrophages and antigens. The fibrotic reaction on the microcapsule surface affects the permeability of the microcapsules and thus prevents insulin, nutrients, and oxygen exchange. Therefore, the fibrotic reaction causes hypoxia and destruction of encapsulated cells, resulting in the necrosis of beta cells and graft failure. Different anti-inflammatory drugs have been analyzed to decrease immune system response, improve biocompatibility, and reduce fibrosis from microcapsules’ surfaces. This strategy enhances the viability of beta cells for a long time (Kim et al., 2021; Samojlik & Stabler, 2021).

The introduction of a drug delivery system into the microcapsules may provide potential local inflammation treatment. Curcumin is a bioactive phytochemical and natural polyphenol present in the rhizome of Curcuma longa (turmeric). During the last decade, many studies have confirmed some attributes of curcumin, such as antitumor, antioxidant, antiviral, and neuroprotective properties (Pehlivanovic, 2020). In addition to these properties, recent studies have confirmed the anti-inflammatory property of curcumin (Menon & Sudheer, 2007). In fact, curcumin can interact with multiple molecular targets involved in inflammation response and minimize the formation of fibrotic cell layers on the implanted microcapsules (Dang et al., 2013). Curcumin decreases the inflammatory reaction by reducing the lipoxygenase’s activity, cyclooxygenase-2 (COX-2), and inducible nitric oxide synthase (iNOS) enzymes; it prevents the generation of the inflammatory cytokines tumor necrosis factor-alpha (TNF-α), migration inhibitory protein, interleukin (IL) −1, −2, −6, −8, and −12, and monocyte chemoattractant protein (MCP) (Hewlings & Kalman, 2017; Jurenka, 2009).

Mass transfer limits the size of any tissue construct because most cells are placed a maximum of 200 μm from the nearby blood-vessel in the body. This space provides adequate diffusion of nutrients and oxygen; consequently, the beta cells can be viable within the body. Thus, the microcapsules’ size is a critical point for encapsulating different cells within biomaterials (Lovett, Lee, Edwards, & Kaplan, 2009). Creating droplets by extrusion through a nozzle is still the primary means for cell encapsulation. This approach is dependent on the different parameters like the rate of the extruded liquid. The more the liquid’s velocity, the smaller the diameter of the microcapsules (Bidoret, Martins, De Smet, & Poncelet, 2017).

Different anti-inflammatory drugs and materials have been studied to enhance the biocompatibility of microcapsules by modulating the immune system’s inflammatory response and subsequently reducing the fibrosis layer on the microcapsules (Kim et al., 2021; Ricci et al., 2005). Kim et al. dissolved Dexamethasone 21-phosphate (Dexa) in 1% chitosan and coated alginate microcapsules with Dexa. They investigated the islets viability, insulin secretion, and fibrosis formation on the microcapsules. They demonstrated that curcumin decreases the fibrotic reaction on the microcapsules (Kim et al., 2021). Ricci et al. assessed the in vitro cytotoxicity of ketoprofen in alginate/poly-l-ornithine/alginate in terms of viability and insulin content. The result indicated that the ketoprofen released from microcapsules decreases inflammatory response (Ricci et al., 2005). Alagpulinsa et al. used an immunomodulatory chemokine, CXCL12, for SC□β cells encapsulating to avoid the formation of fibrotic response. CXCL12 caused glycaemia for a long period by removing fibrotic reaction (Alagpulinsa et al., 2019). Dang et al. co-encapsulated 16 anti-inflammatory drugs and islet into microcapsules created by extrusion of the islet-alginate suspension at a low flow rate and 6 kV voltage and injected them subcutaneously to evaluate the immunosuppressive effects. Among selected drugs, curcumin and Dexa were recognized as the most effective anti-inflammatory drugs by lowering the inflammatory proteases’ activity and decreasing reactive oxygen species (Dang et al., 2013).

This study aims to co-encapsulate beta cells and curcumin within the micro-size particles as an anti-inflammatory drug to investigate the impact of curcumin on insulin release and cell viability. For creating microcapsules, the velocity of the syringe pump is optimized to form a jet-breaking regime in which the solution exits the needle like a jet and breaks-up in droplets form. Our study involves the in vitro characterization of the carrying microcapsule and assessment of drug release, cell viability, and beta cell function.

## 3. Materials and Methods

### Materials

RIN-5F cells were purchased from the Tehran’s Iranian Biological Resource Center. Calcium chloride was obtained from Merck (Darmstadt, Germany). Medium viscosity sodium alginate, Krebs-Ringer Bicarbonate Buffer (With 1800 mg/L glucose, without calcium chloride), MTT solution (3-(4,5-Dimethylthiazol-2-yl)-2,5-Diphenyltetrazolium Bromide), glucose, curcumin, sodium bicarbonate, and Phosphate buffered saline were obtained from Sigma Aldrich (St Louis, MO, USA). RPMI 1640 medium and fetal bovine serum (FBS) were obtained from GIBCO (Carlsbad, CA, USA). 1% penicillin/streptomycin, Trypsin/EDTA, and DMSO were achieved from BioIdea, Iran.

### Cell Culture

The pancreatic beta cells (**RIN**-**5F cell line**) were cultured in a 25 cm^3^ culture flask that contained RPMI 1640 medium with 10% FBS and 1% penesterep antibiotic. Flask was placed at 37°C and 5% CO_2_ incubator with 95% humidity. After cells reached proper growth, sub-culturing was done.

### Cell Microencapsulation

Medium viscosity alginate was used to produce microcapsules. At first, alginate was dissolved in sterile water to prepare 1.5% (w/v) alginate solution. 100 mM Calcium chloride solution was used as a cross-linking solution. Alginate and calcium chloride solutions were sterilized with a 0.22 μm syringe filter. Sterile alginate solution was mixed with curcumin at 1.0 mg/ml and stirred to confirm that the curcumin was homogenously dispersed in alginate solution. The beta cells were detached from the culture flask by adding 2 ml trypsin and incubating for 2 minutes. After 2 minutes, 4 ml RPMI was added and centrifuged with 1500 rpm for 5 minutes. For encapsulating the beta cells within microcapsules, as shown in figure 1, alginate solution with or without curcumin was mixed with beta cells (2×10^4^ cells/ml). Then a 30 gauge needle and syringe pump were used to produce microcapsules. The rate of the syringe pump was adjusted to 5, 10, 15, 20, and 25 ml/min to find a jet-breaking regime that could reduce the diameter of the microcapsules and produce micro-size particles. The syringe was filled with the alginate solution that contained cells and the drug. Then the syringe pump was adjusted on top of the calcium chloride bath. The microcapsules were produced when the liquid exited the needle like a jet regime and broke-up as droplets. Then the formed microcapsules were incubated in a cross-linking solution (100mM CaCl_2_) for 5 minutes. After gelation, the microcapsules were rinsed twice with PBS to remove any extra calcium ions from the microcapsules. After washing, the microcapsules were moved to RPMI media and incubated at 37 °C, with 95% humidity and 5% CO_2_.

**Figure 1.**
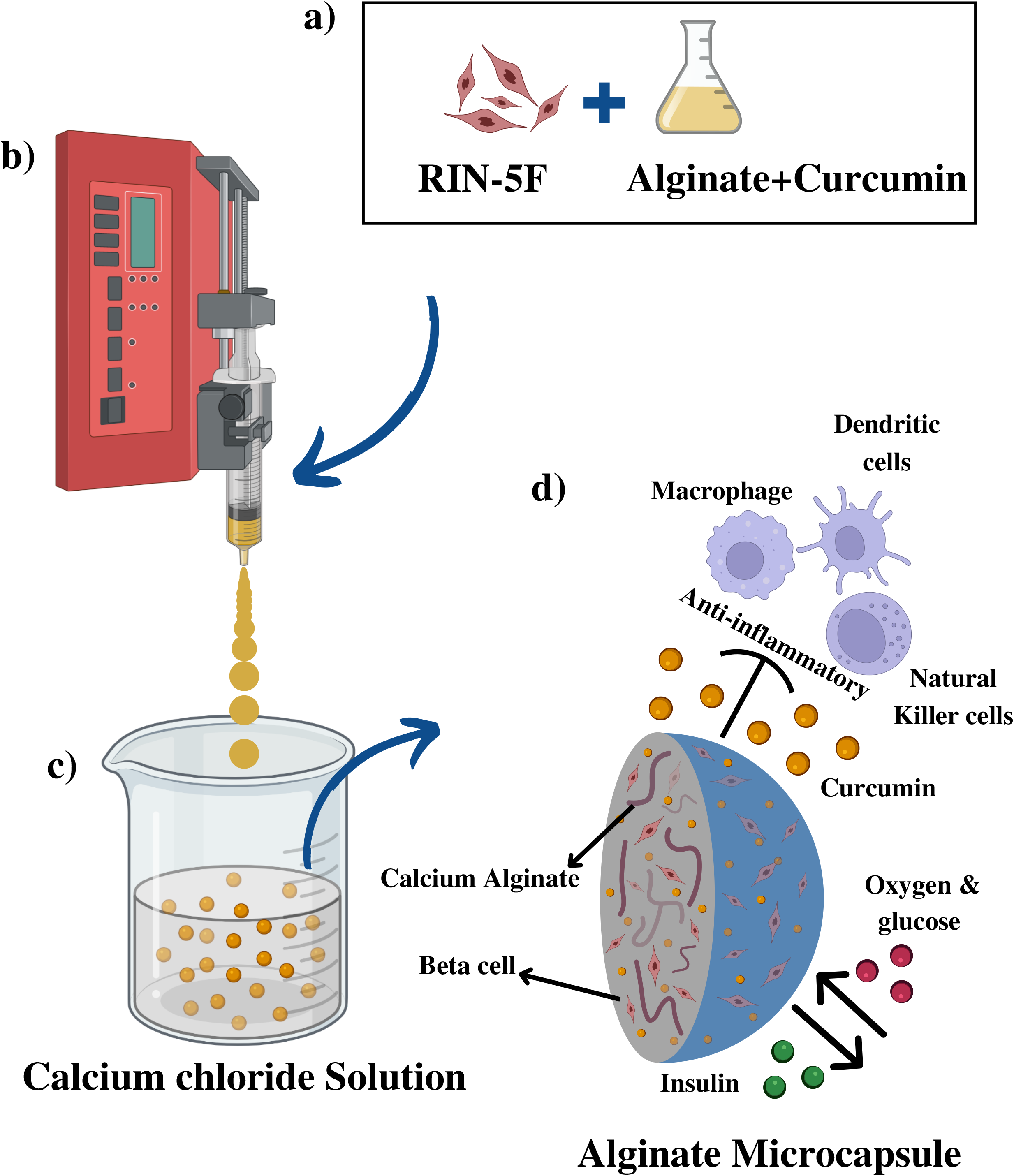
Schematic of the co-encapsulation of RIN-5F with curcumin using the jet-breaking technic. (a) Rin-5F cell suspension in the alginate-curcumin solution. (b) Droplet formation by jet-breaking technic. (c) Cross-linking solution (100 mM calcium chloride). (d) Co-encapsulated cells and curcumin within alginate microcapsules, which plays an anti-inflammatory drug and blocks the immune system’s cells.

### Particle size distribution

To evaluate the diameter of the microparticles, which is an essential factor in cell encapsulation for oxygen and nutrient diffusion, 200±3 microcapsules were selected randomly. Microcapsules’ photos were captured with a light microscope, then the average diameter of microcapsules was measured by Image J software.

### Stability of microcapsules

Microparticles were prepared and randomly selected (n = 200±5) and placed in the 3 ml PBS then incubated at 37 °C. The microcapsules were observed under a light microscope after 1, 3, and 7 days. The number of damaged and broken microparticles was recorded (Nikravesh et al., 2017).

### Swelling ratio of microcapsules

After the microcapsules were generated, they were freeze-dried and the diameter was recorded by light microscope. Then the microcapsules were soaked in deionized water at 37 °C. After 10, 20, 30, 60 and 120 minutes, the diameter of microcapsules was measured after removing the excess fluid (Roshanbinfar & Salahshour Kordestani, 2013). The swelling ratio was computed by the formulation (1) given by Liu et al (Liu et al., 2004).

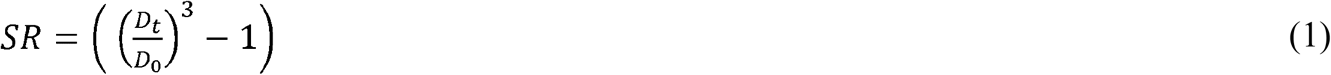

where D_0_ and D_t_ represent the diameter of the freeze-dried microcapsules before and after swelling, respectively.

### In vitro curcumin cytotoxicity

Cytotoxicity assays were performed to investigate curcumin’s toxic effect on beta cells. Briefly, three experimental conditions were examined. Free cells and alginate microcapsules were used as control, and curcumin-alginate microcapsules were used as the treated group.

All groups were maintained in RPMI medium (Ricci et al., 2005). Viability was assessed by MTT-assay. This test demonstrates proliferation grade and cell living situation. This assay was assessed by the amount of insoluble, purple formazan crystals produced by mitochondrial dehydrogenase enzymes in viable cells. Each group of microcapsules and free cells containing 6000 beta cells were incubated in 500 μl of culture media and 50 μl of MTT solution (0.5 mg/ml) for 24 hours in a 24-well plate. After 24 hours, 500 μl of DMSO was added to each wall, and the well plate was wrapped in aluminum foil and incubated for 15 minutes. After incubation for 15 minutes, the formazan crystals were quantified by measuring the absorbance of each wall using an ELIZA reader at a wavelength of 490 nm (Haque et al., 2005). Cytotoxicity activity was calculated according to the formulation (2) equation:

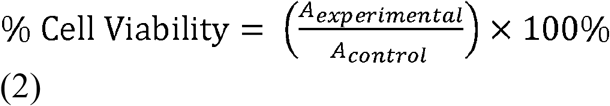

where, A_control_ represents absorbance of free cells and A_experimental_ is absorbance of encapsulated cells (Sookkasem, Chatpun, Yuenyongsawad, & Wiwattanapatapee, 2015).

### Glucose-Stimulated Insulin Secretion (GSIS) assay

To investigate the RIN-5F glucose response, GSIS assay was completed 1, 3, and 7 days after cell encapsulation. Similar to the cytotoxicity test, three groups (free cells, alginate microcapsules, and curcumin-alginate microcapsules) were studied for the GSIS assay. Free cells and microcapsules were rinsed and placed in Krebs-Ringer bicarbonate (KRB) and incubated for 30 minutes. Then, KRB was removed and replaced with KRB having 3.3 mM glucose and placed in CO_2_ incubator for 2 hours. Then, supernatant of each sample were gathered to measure insulin secretion, and microcapsules were rinsed and placed in incubator for 2 hours in KRB holding 16.6 mM glucose. At the end, supernatants were gathered to quantify insulin secretion in response to increasing glucose. The insulin of each sample was measured with the C-Peptide & Insulin AccuBind VAST ELISA Kits (Monobind Inc., USA). Three independent experiments were performed for each group. (Espona-Noguera et al., 2018).

### In vitro curcumin release test

To investigate the curcumin release from microcapsules, the microcapsules containing curcumin were immersed in a certain amount of buffer phosphate solution and placed in an incubator at 37 °C. Then, at certain hours, each sample’s phosphate buffer medium was gathered and centrifuged at 1500 rpm for 5 minutes and absorbance was measured by the spectrophotometer at 420 nm.

### Statistical analysis

All experiments were completed in three times, and data were represented as mean ± standard deviation. One-way ANOVA was done with Microsoft Excel for Statistical analysis, and the significant level was selected as P < 0.05.

## RESULT

### Characterization of microcapsules

The diameter of the microcapsules can be altered by changing the alginate-cell suspension’s rate, space between the needle and cross-linker solution bath, and the needle gauge. In this study, the rate of the alginate was altered to achieve micro-sized capsules. The result indicated that the maximum syringe pump rate produced micro-size capsules with uniform shape (Figure 2 a and b). The optimized parameters used were syringe size of 10 ml, syringe diameter of 14 mm, 30 gauge needle, and height between the needle and cross-linker solution (CaCl_2_) was 5.0 cm roughly.

**Figure 2.**
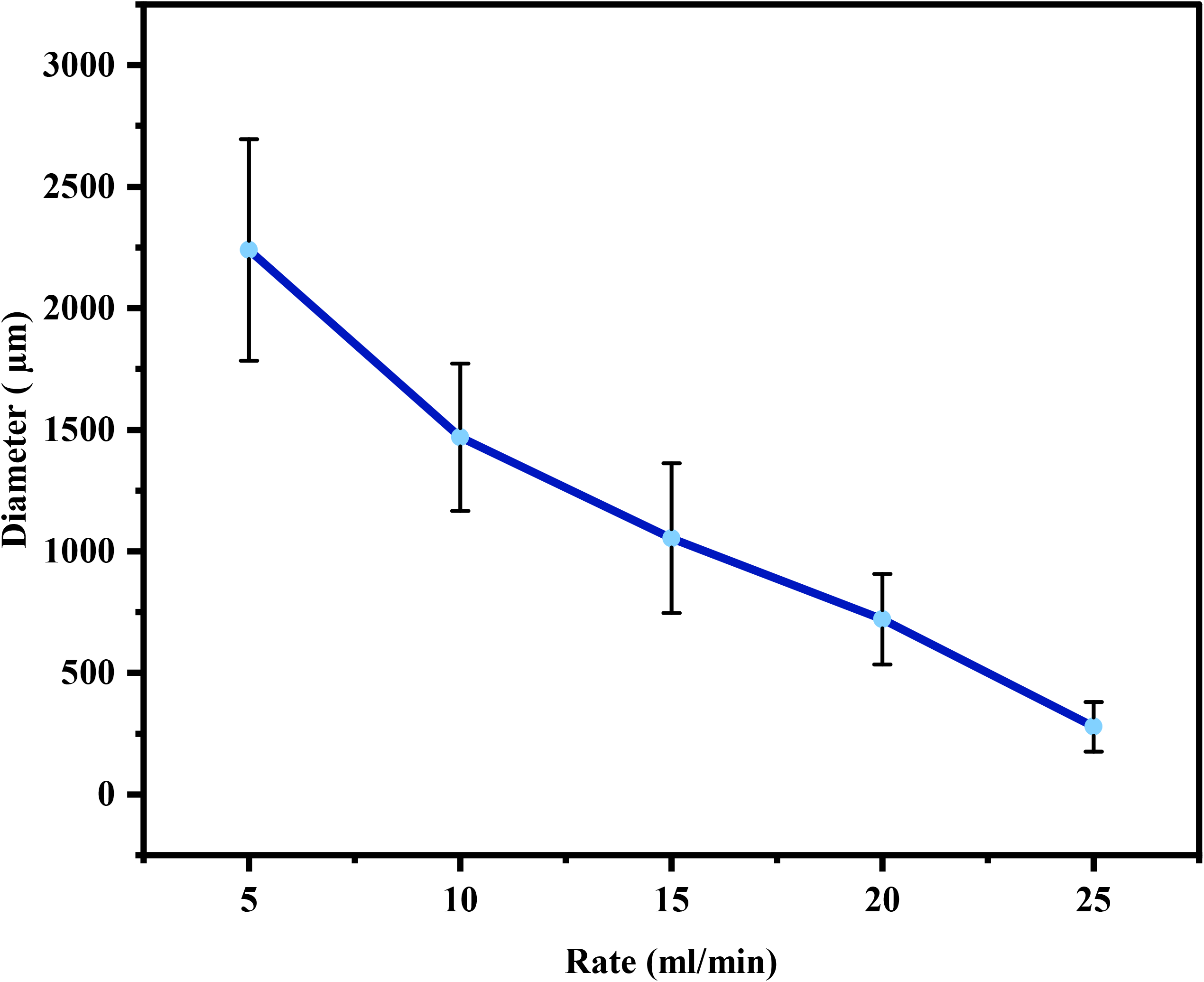
Optimization of syringe pump’s flow rate. (a) Optimization of syringe pump’s flow rate for forming micro-size capsules (n = 100 ±5 capsules).

The radius of microcapsules is an essential factor in cell encapsulation, and it should be less than 200 μm to deliver sufficient oxygen and nutrients. In this study, the particle-size distribution of the alginate microcapsules with cells ranged from 50 to 500 μm, and the average diameter was 288 μm (see Figure 2 c). The results indicated that the average radius of microcapsules was less than 200 μm. Thus, the cells can be viable within the microcapsules. The diameter of microcapsules placed in the range of 200–300 μm and followed the Gaussian distribution. The maximum number of microcapsules’ diameter placed in the range of 250-300 μm (Figure 2 c).

### Stability of microcapsules

Alginate microcapsules should have long stability to apply for long-term cell encapsulation. Breaking microcapsules can change product and drug release kinetics and destroy the encapsulated cells by the immune system (Moya, Morley, Khanna, Opara, & Brey, 2012). Approximately 200±5 alginate microcapsules were observed by a light microscope to find ruptured microcapsules for 7 days. Only 5 damaged microcapsules were observed. Thus, more than 97% of microcapsules remained intact during 7 days (Figure 3 c and d). That means the microcapsules were mechanically stable enough and could protect the encapsulated beta cells over a long time. The stability of microcapsules is agree with Nikravesh’s study (Nikravesh et al., 2017).

**Figure 3.**
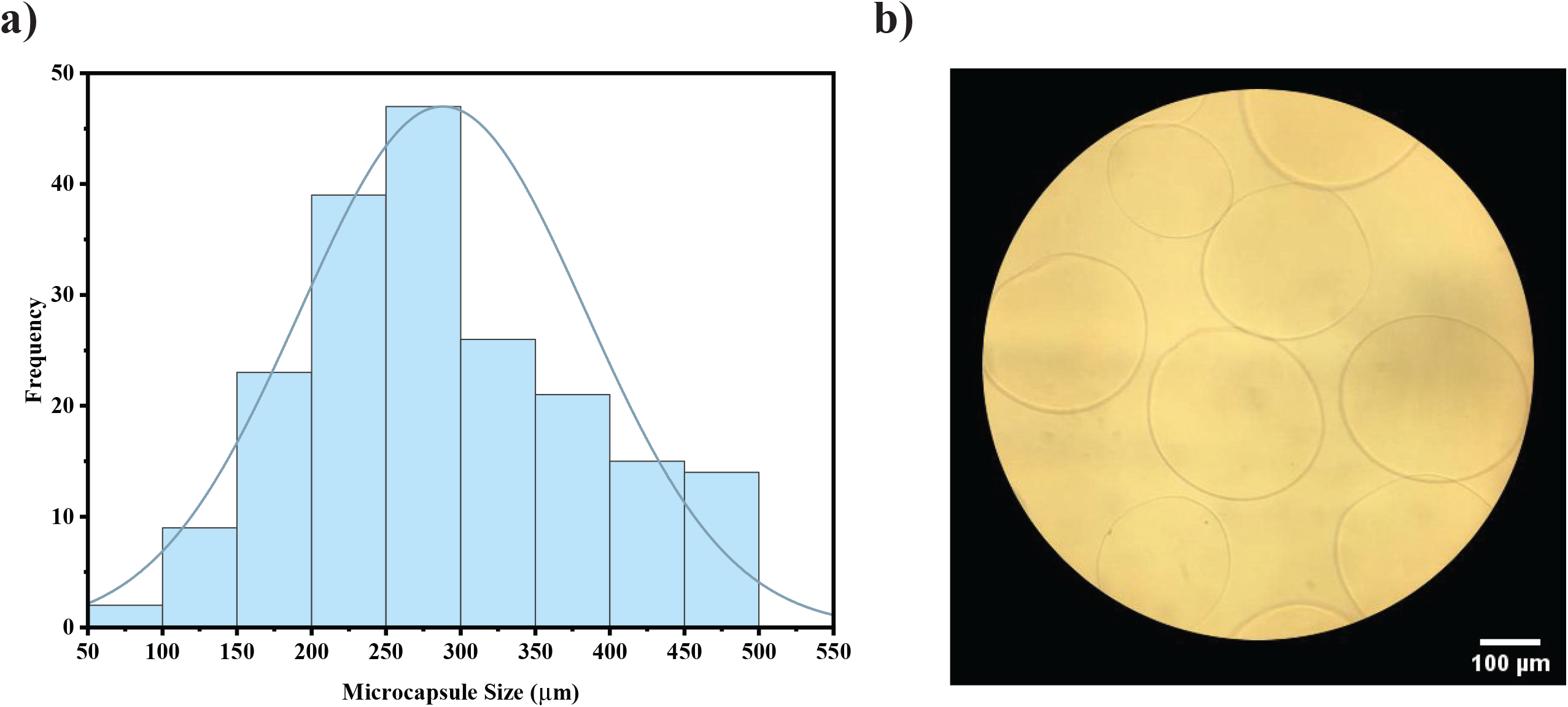
(a) Light microscopic image of optimized cell-free microcapsules (Alginate concentration = 1.5%, CaCl2 concentration = 100 mM, pump flow rate = 25 ml/min, and needle size = 30 created. (b) Histogram of diameter distribution of alginate microcapsules created by jet-breaking technic (n=200±3 Microcapsules). Scale bar = 100 μm.

### Swelling ratio of microcapsules

The swelling ratio of microcapsules represents the capsules’ capacity to absorb body fluid and transfer nutrients and metabolites. Therefore, this is an essential property of hydrogel-based biomaterials (Sarker et al., 2014). Samples were placed in deionized water for 120 minutes, and a dimensional assay was done in the point times with a light microscope (Figure 3 a). At the end of the 120 minutes, the average diameter of microcapsules was increased by approximately 250 μm, and the swelling ratio was increased 12-fold (Figure 3 b). Thus, the alginate microcapsules could keep and diffuse the deionized water. The swelling results of alginate microcapsules agree with the results reported by Sarker et al. (Sarker et al., 2014).

### In vitro curcumin cytotoxicity

The viability of encapsulated beta cells with or without curcumin was analyzed by MTT assay 1, 3, and 7 days after the encapsulation process and compared with free cells’ viability. MTT assay was completed to assess the possible toxic impact of curcumin and alginate membrane on beta cells. Images captured with a light microscope on day 7 from encapsulated cells with and without curcumin showed that beta cells morphology remained undamaged (Figure 4 a and b). During the 7 days (Figure 4 c), it is evident from the absorbance chart that cell viability in the microcapsules enhances with time. The absorbance of the encapsulated cells within alginate-curcumin increased from 0.163 to 0.832 after 7 days of incubation. 7 days after microencapsulation, the percentage of viable cells in the alginate-curcumin group (90.34%±1.31%) was analogous to that of the alginate group (91.97%±1.06%) and did not differ from that of the free cells (100%). Thus, the alginate membrane and curcumin concentration employed did not induce toxic impacts on the viability of beta cells and can support proliferation. The results proved that the encapsulated cells in alginate had good viability, similar to those in Duruksu’s report (Duruksu et al., 2018).

**Figure 4.**
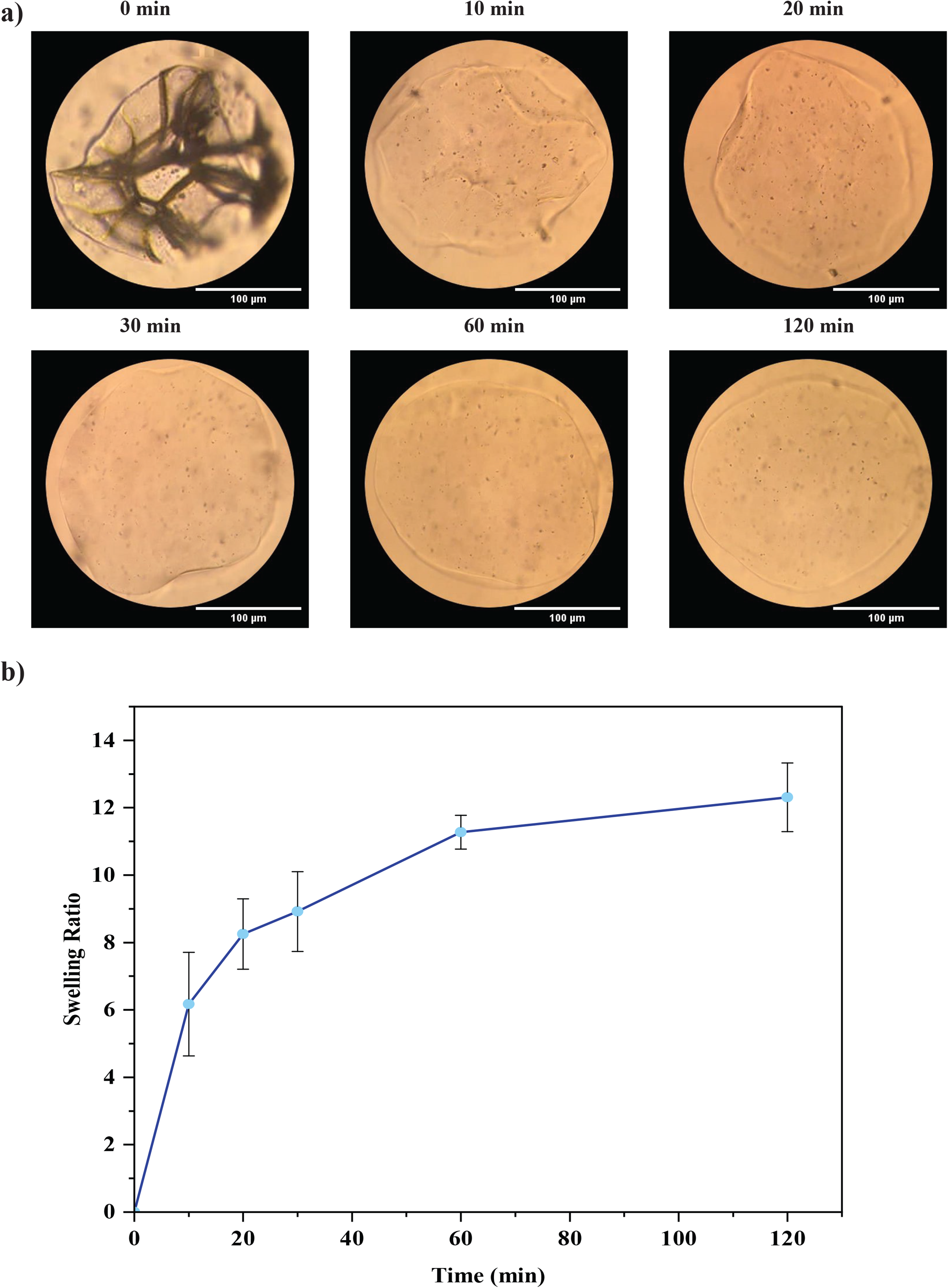
Characterization of alginate microcapsules. (a) Optical microscopy image of blank microcapsule after freeze-drying (0 min), 10 min, 20 min, 30 min, 60 min, and 120 min after placing in PBS (n=100 microcapsules). (b) The swelling ratio of alginate microcapsules placed in PBS at pH=7.4 at different times (n=100 microcapsules).

### Insulin secretion in response to glucose

In response to increasing glucose concentration in the cell culture, the insulin secretion level, which is vital to investigate the functionality of microencapsulated beta cells, was assessed using the GSIS (Glucose-Stimulated insulin Secretion) assay in vitro. Figure 5 a-c reveal that encapsulated RIN-5F cells within 1.5% alginate microcapsules revealed a considerable increase in secreted insulin when glucose was enhanced from 3.3 mM to 16.6 mM (P < 0.05). On day 1, free cells, encapsulated cells without, and with curcumin secreted 1.53 μIU/mL ± 0.38, 1.49 μIU/mL ± 0.23, and 1.46 μIU/mL ± 0.14 in response to 3.3 mM and 34.09 μIU/mL ± 7.57, 33.33 μIU/mL ± 10.10, and 42.39 μIU/mL ± 19.78 in response to 16.6 mM glucose concentration, respectively. The insulin secretion levels in encapsulated cells were not significantly different from free cells (P > 0.05). The same trends were observed on days 3 and 7 after encapsulation. The results indicated no significant difference between insulin concentration from the encapsulated and free cells, meaning that the alginate membranes have semi-permeability, and the alginate membrane and curcumin did not influence beta cell function. In addition, we found a significant difference in the quantity of insulin secreted during the entire study time, and insulin secretion had an upward trend until the seventh day (Figure 5 d), which is similar to the value noted elsewhere (Nikravesh et al., 2017). This result indicated the proliferation of cells within the microcapsules (P <0.05). Consequently, curcumin did not affect the survival and insulin-secreting ability of beta cells.

**Figure 5.**
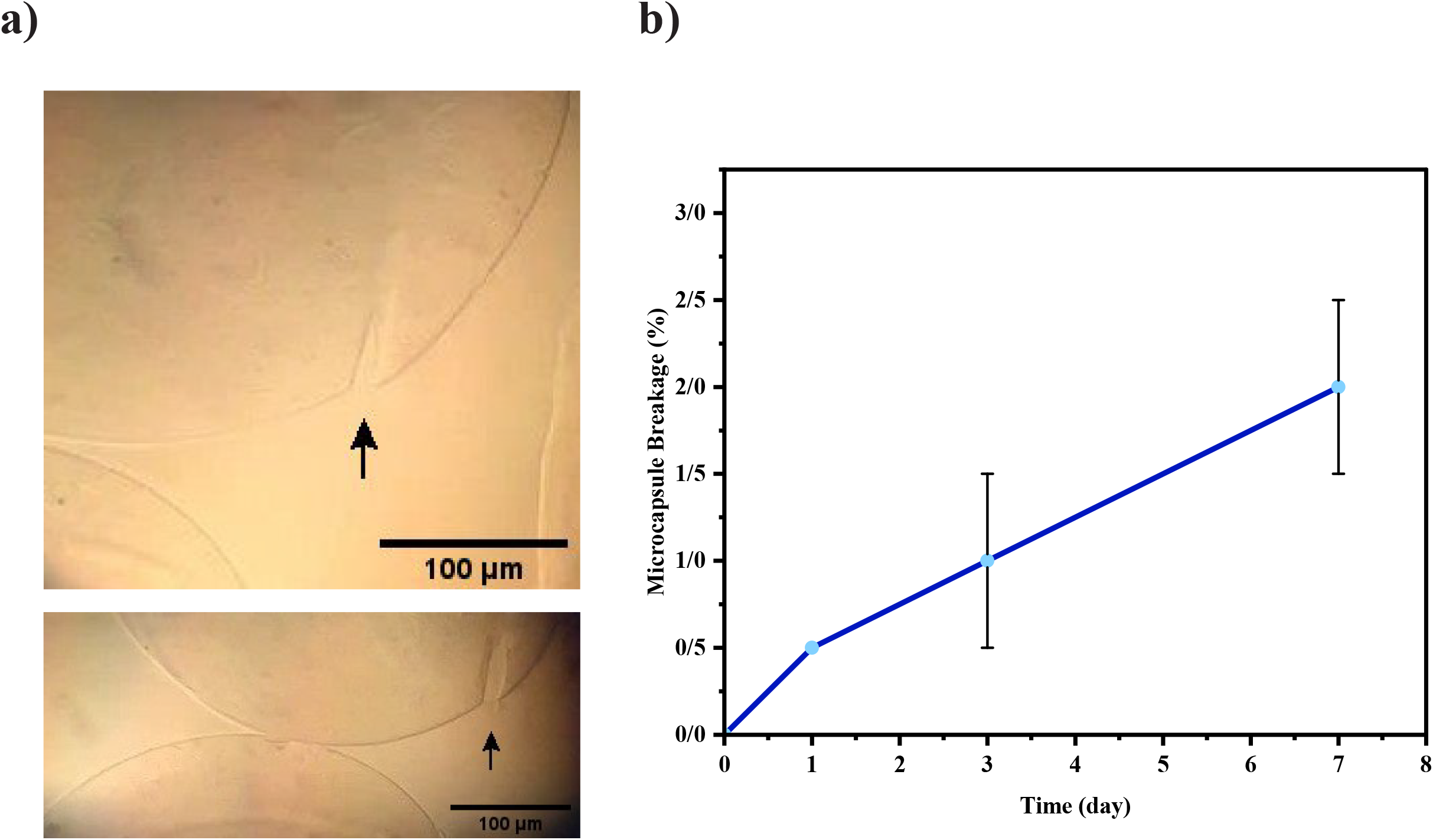
(a) Microscopy image of damaged microcapsules. (d) Percent of breakage microcapsules during 7 days (n=200 microcapsules). Scale bar = 100 μm.

### In vitro curcumin release test

According to the results, the release rate of curcumin from alginate microcapsules is about 0.26% (Figure 6), which is probably because of the less aqueous solubility of curcumin, which leads to a slower release of curcumin. Accordingly, curcumin remains in alginate microcapsules for a prolonged period of time. Therefore, its efficacy in lowering the fibrotic response will be longer and keep the cells viable. Remaining the curcumin in the microcapsules for long time and releasing slowly is correlate with Dang et al.’s observation (Dang et al., 2013).

**Figure 6.**
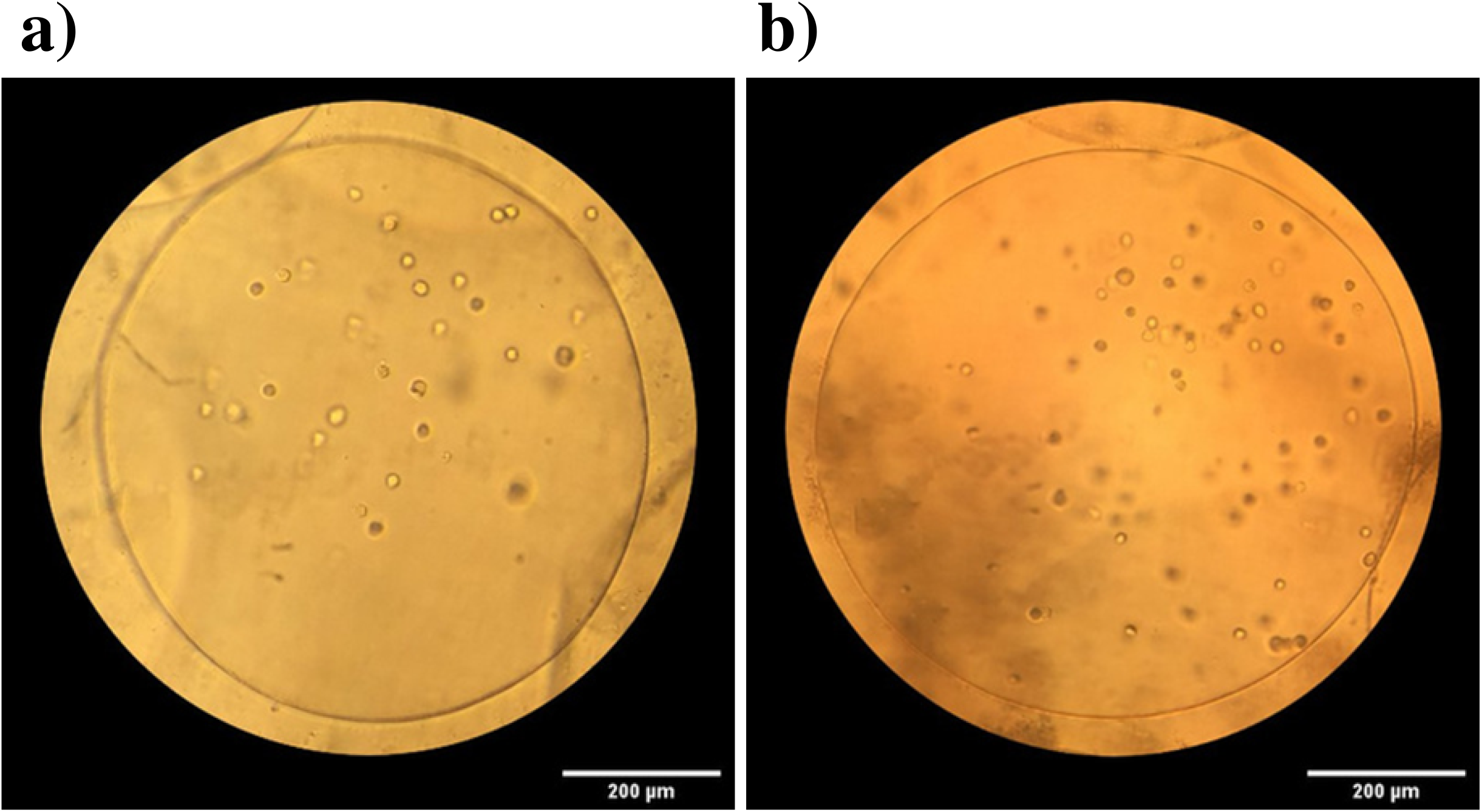
Microscopic image of encapsulated cells (a) without curcumin and (b) with curcumin.

**Figure 7.**
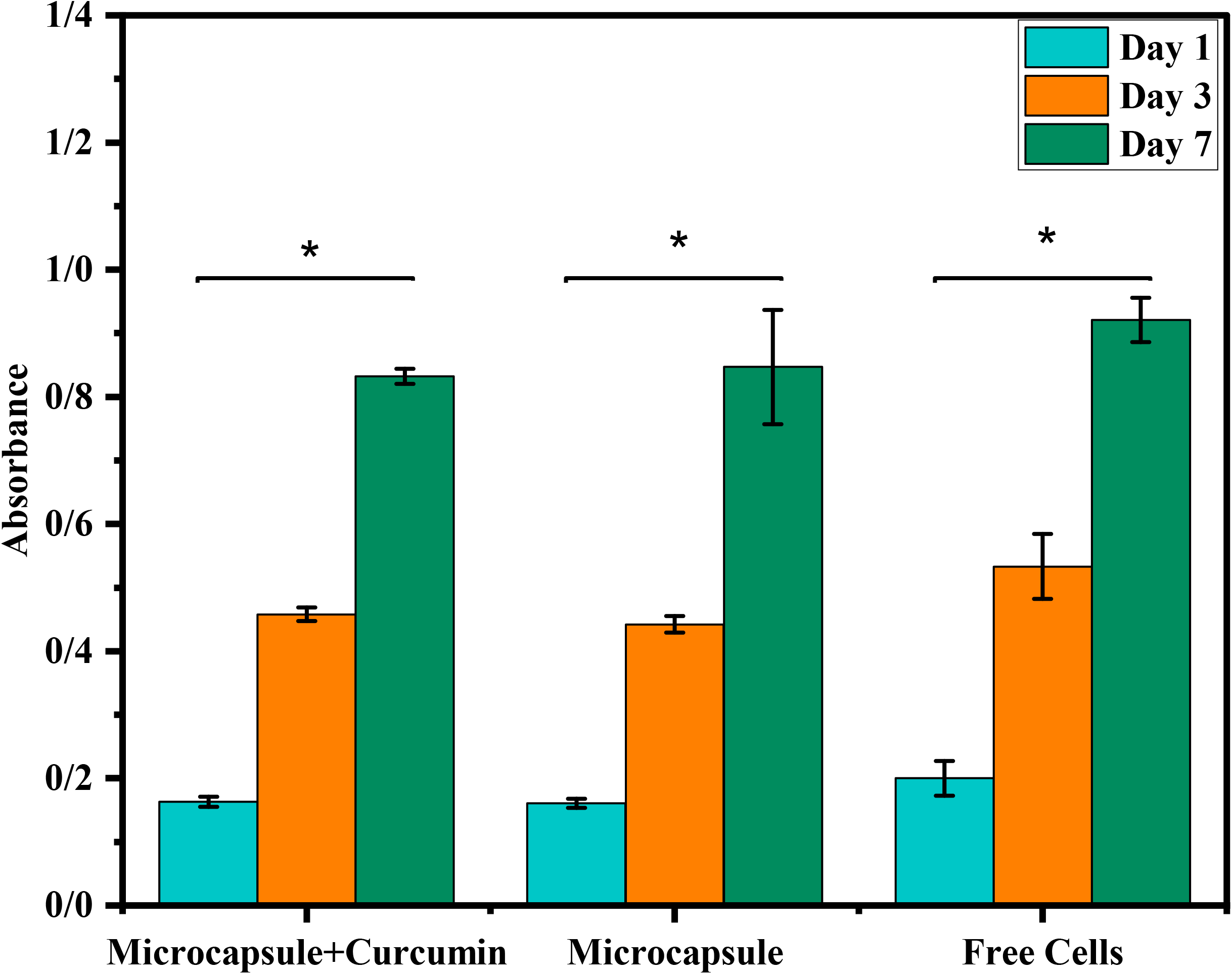
Comparison of cell viability between free cells, encapsulated cells, and co-encapsulated cells with curcumin by MTT assay. The absorbance of different groups 1, 3, and 7 days after encapsulation (n=3). ^*^p<0.05. Scale bar = 200 μm.

**Figure 8.**
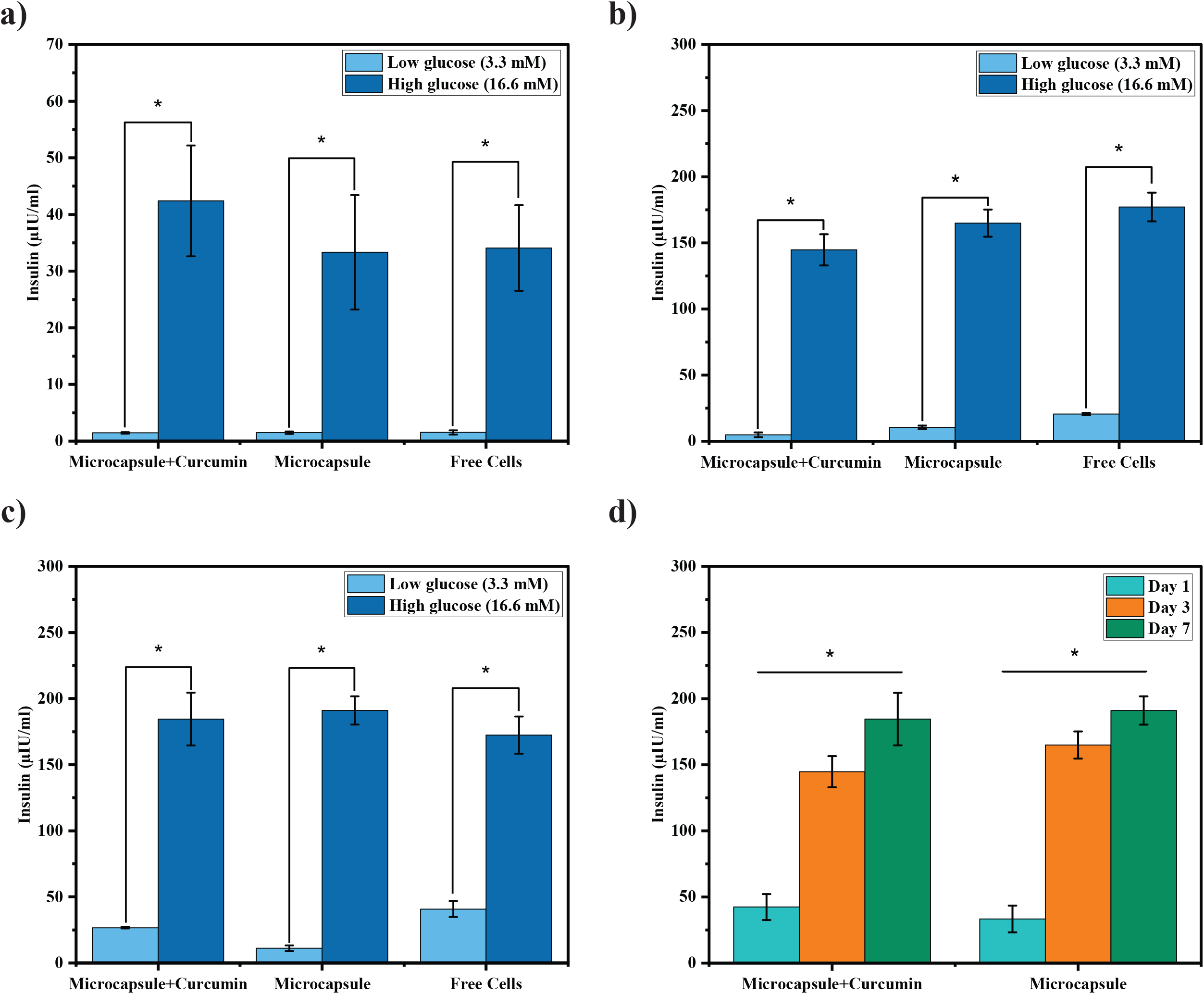
Insulin secretion levels are determined by ELISA. (a) The secreted insulin in response to the low-glucose (3.3 mM) and high-glucose (16.6 mM) concentration from free cells, encapsulated cells, and co-encapsulated cells with curcumin on day 1, (b) day 3, (c) day 7. (d) Comparison of secreted insulin from encapsulated cells with and without curcumin on different days. n=3, *p<0.05.

**Figure 9.**
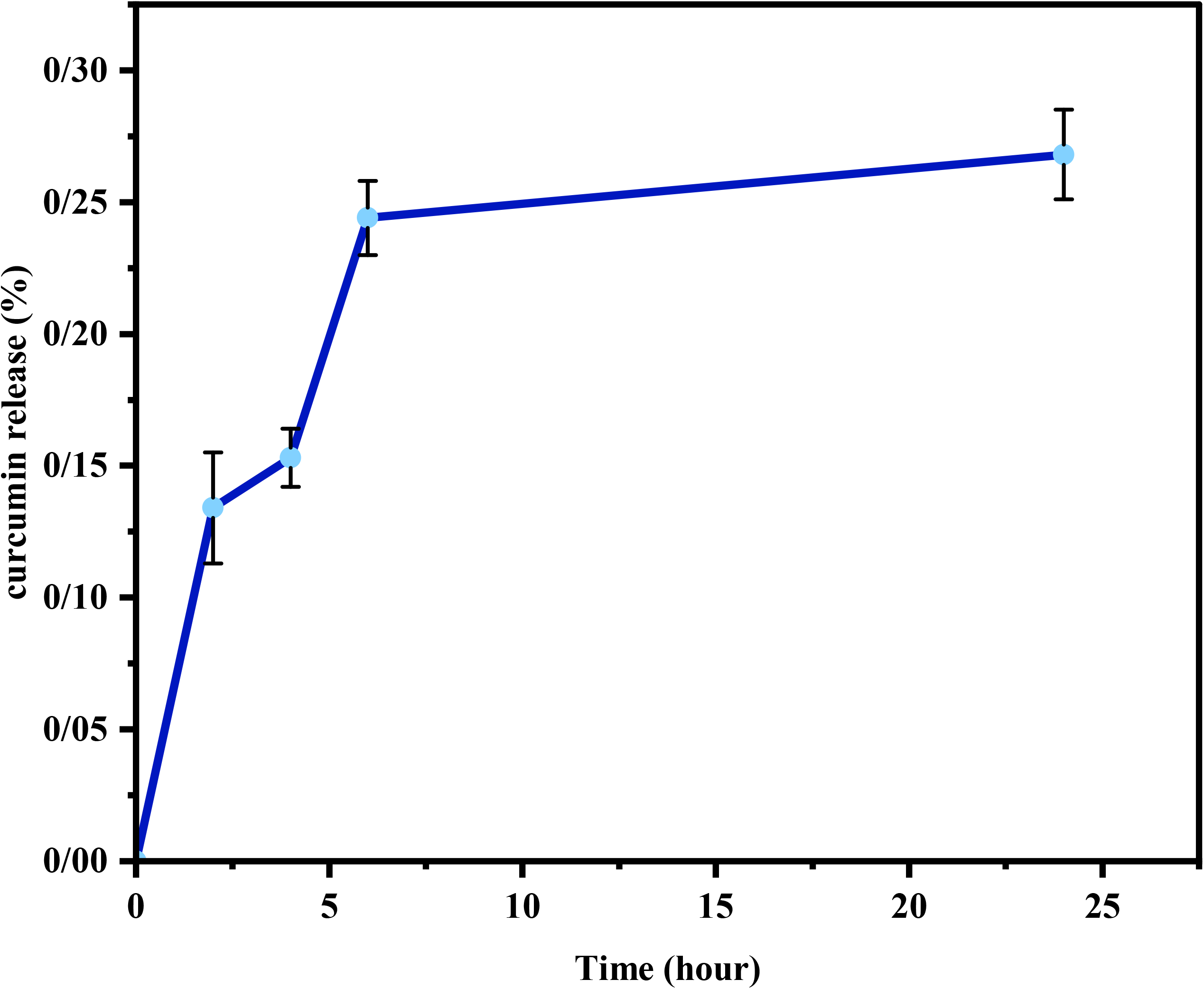
Percent of curcumin released from microcapsules during 24 hours (n=3).

## CONCLUSION

In this study, the jet-breaking technic and alginate were used to create microencapsulated beta cells with and without curcumin. This technic provides micro-size capsules with a uniform shape in a short time. The radius of microcapsules was less than 200 μm, which provided nutrients for encapsulated cells and prevented the death of the cells. The free beta cells were used as a control in this study. Our result indicated that the beta cells’ viability and consequently the insulin level in media were not significantly different from free cells. Thus, the curcumin had no adverse effect on the cell function and survival rate. In addition, the cell viability and insulin levels were significantly different on different days, which implied the cell proliferation in the microcapsules. Our results suggested that the combined use of anti-inflammatory drugs and beta cells within microcapsules is a promising method for treating type I diabetes.

## AUTHOR CONTRIBUTIONS

**Zahra Hosseinzadeh:** Conceptualization, Methodology; Software; Formal Analysis; Investigation; Data Curation; Writing – Original Draft. **Iran Alemzadeh:** Investigation; Writing – Review and Editing, Project Administration, Funding Acquisition, Supervision. **Manouchehr Vossoughi:** Investigation; Writing – Review and Editing, Project Administration, Funding Acquisition, Supervision.

## ACKNOWLEDGMENTS

The authors thankfully acknowledge the experimental support by Biochemical and Bioenvironmental Engineering Research Center (BBRC) and the Chemical and Petroleum Engineering Department at Sharif University of Technology.

## ETHICAL STATEMENT

The authors confirm that the ethical policies of the journal, as noted on the journal’s author guidelines page, have been adhered to and the appropriate ethical review committee approval has been received. The study conformed to the US Federal Policy for the Protection of Human Subjects.

## REFERENCES

Alagpulinsa, D. A., Cao, J. J., Driscoll, R. K., Sîrbulescu, R. F., Penson, M. F., Sremac, M., Melton, D. A. (2019). Alginate-microencapsulation of human stem cell–derived β cells with CXCL 12 prolongs their survival and function in immunocompetent mice without systemic immunosuppression. American Journal of Transplantation, 19(7), 1930–1940. doi:https://doi.org/10.1111/ajt.15308

Atkinson, M. A.-O., Campbell-Thompson, M. A.-O., Kusmartseva, I. A.-O., & Kaestner, K. A.-O. X. (2020). Organisation of the human pancreas in health and in diabetes. (1432-0428 (Electronic)). doi:https://doi.org/10.1007/s00125-020-05203-7

Bansal, A., D’Sa, S., & D’Souza, M. J. (2019). Biofabrication of microcapsules encapsulating beta-TC-6 cells via scalable device and in-vivo evaluation in type 1 diabetic mice. International Journal of Pharmaceutics, 572, 118830. doi:https://doi.org/10.1016/j.ijpharm.2019.118830

Bidoret, A., Martins, E., De Smet, B. P., & Poncelet, D. (2017). Cell Microencapsulation: Dripping Methods. (1940-6029 (Electronic)). doi:https://doi.org/10.1007/978-1-4939-6364-5_3

Chan, K. H., Krishnan, R., Alexander, M., & Lakey, J. R. (2017). Developing a Rapid Algorithm to Enable Rapid Characterization of Alginate Microcapsules. Cell transplantation, 26(5), 765–772. doi:https://doi.org/10.3727/096368916X693446

Christoffersson, G. (2022). Towards effective beta-cell replacement through understanding and targeting of the autoreactive immune response during onset of type 1 diabetes. Journal of Immunology and Regenerative Medicine, 15, 100057. doi:https://doi.org/10.1016/j.regen.2021.100057

Dang, T. T., Thai, A. V., Cohen, J., Slosberg, J. E., Siniakowicz, K., Doloff, J. C., Gu, Z. (2013). Enhanced function of immuno-isolated islets in diabetes therapy by co-encapsulation with an anti-inflammatory drug. Biomaterials, 34(23), 5792–5801. doi:https://doi.org/10.1016/j.biomaterials.2013.04.016

de Vos, P., Faas, M. M., Strand, B., & Calafiore, R. (2006). Alginate-based microcapsules for immunoisolation of pancreatic islets. Biomaterials, 27(32), 5603–5617. doi:https://doi.org/10.1016/j.biomaterials.2006.07.010

Duruksu, G., Polat, S., Kayiş, L., Gürcan, N. E., Gacar, G., & Yazir, Y. (2018). Improvement of the insulin secretion from beta cells encapsulated in alginate/poly-L-histidine/alginate microbeads by platelet-rich plasma. Turkish Journal of Biology, 42(4), 297–306. doi:https://doi.org/10.3906/biy-1802-13

Espona-Noguera, A., Ciriza, J., Cañibano-Hernández, A., Fernandez, L., Ochoa, I., Del Burgo, L. S., & Pedraz, J. L. (2018). Tunable injectable alginate-based hydrogel for cell therapy in Type 1 Diabetes Mellitus. International journal of biological macromolecules, 107, 1261–1269. doi:http://dx.doi.org/10.1016/j.ijbiomac.2017.09.103

Espona-Noguera, A., Ciriza, J., Cañibano-Hernández, A., Orive, G., Hernández, R. M., Saenz del Burgo, L., & Pedraz, J. L. (2019). Review of advanced hydrogel-based cell encapsulation systems for insulin delivery in type 1 diabetes mellitus. Pharmaceutics, 11(11), 597. doi:https://doi.org/10.3390/pharmaceutics11110597

Ghoneim, M. A., Refaie, A. F., Elbassiouny, B. L., Gabr, M. M., & Zakaria, M. M. (2020). From mesenchymal stromal/stem cells to insulin-producing cells: progress and challenges. Stem Cell Reviews and Reports, 16(6), 1156–1172. doi:https://doi.org/10.1007/s12015-020-10036-3

Haque, T., Chen, H., Ouyang, W., Martoni, C., Lawuyi, B., Urbanska, A., & Prakash, S. (2005). Investigation of a new microcapsule membrane combining alginate, chitosan, polyethylene glycol and poly-L-lysine for cell transplantation applications. The International journal of artificial organs, 28(6), 631–637. doi:https://doi.org/10.1177/039139880502800612

Hewlings, S. J., & Kalman, D. S. (2017). Curcumin: A review of its effects on human health. Foods, 6(10), 92. doi:https://doi.org/10.3390%2Ffoods6100092

Holt, E. H., Lupsa, B., Lee, G. S., Bassyouni, H., & Peery, H. E. (2022). Chapter 7 - The pancreatic islets. In E. H. Holt, B. Lupsa, G. S. Lee, H. Bassyouni, & H. E. Peery (Eds.), Goodman’s Basic Medical Endocrinology (Fifth Edition) (pp. 203–237): Elsevier.

Jara, C., Oyarzun-Ampuero, F., Carrión, F., González-Echeverría, E., Cappelli, C., & Caviedes, P. A.-O. (2020). Microencapsulation of cellular aggregates composed of differentiated insulin and glucagon-producing cells from human mesenchymal stem cells derived from adipose tissue. (1758-5996 (Print)). doi:https://doi.org/10.1186/s13098-020-00573-9

Jurenka, J. S. (2009). Anti-inflammatory properties of curcumin, a major constituent of Curcuma longa: a review of preclinical and clinical research. Alternative medicine review, 14(2). Retrieved from https://www.researchgate.net/deref/ www.ncbi.nlm.nih.gov%2Fpubmed%2F19594223

Kim, M. J., Park, H.-S., Kim, J.-W., Lee, E.-Y., Rhee, M., You, Y.-H., Yoon, K.-H. (2021). Suppression of fibrotic reactions of chitosan-alginate microcapsules containing porcine islets by dexamethasone surface coating. Endocrinology and Metabolism, 36(1), 146. doi:https://doi.org/10.3803/enm.2021.879

Li, W., Zhao, R., Liu, J., Tian, M., Lu, Y., He, T., Chen, L. (2014). Small islets transplantation superiority to large ones: implications from islet microcirculation and revascularization. (2314-6753 (Electronic)). doi:https://doi.org/10.1155/2014/192093

Liu, X., Xue, W., Liu, Q., Yu, W., Fu, Y., Xiong, X., Yuan, Q. (2004). Swelling behaviour of alginate–chitosan microcapsules prepared by external gelation or internal gelation technology. Carbohydrate Polymers, 56(4), 459–464. doi:https://doi.org/10.1016/j.carbpol.2004.03.011

Lovett, M., Lee, K., Edwards, A., & Kaplan, D. L. (2009). Vascularization strategies for tissue engineering. Tissue Engineering Part B: Reviews, 15(3), 353–370. doi:https://doi.org/10.1089%2Ften.teb.2009.0085

Menon, V. P., & Sudheer, A. R. (2007). Antioxidant and anti-inflammatory properties of curcumin. The molecular targets and therapeutic uses of curcumin in health and disease, 105–125. doi:https://doi.org/10.1007/978-0-387-46401-5_3

Mochizuki, Y., Kogawa, R. A.-O., Takegami, R., Nakamura, K. A.-O., Wakabayashi, A., Ito, T., & Yoshioka, Y. (2020). Co-Microencapsulation of Islets and MSC CellSaics, Mosaic-Like Aggregates of MSCs and Recombinant Peptide Pieces, and Therapeutic Effects of Their Subcutaneous Transplantation on Diabetes. LID - 10.3390/biomedicines8090318 [doi] LID - 318. (2227-9059 (Print)). doi:https://doi.org/10.3390/biomedicines8090318

Moya, M. L., Morley, M., Khanna, O., Opara, E. C., & Brey, E. M. (2012). Stability of alginate microbead properties in vitro. Journal of Materials Science: Materials in Medicine, 23(4), 903–912. doi:https://doi.org/10.1007/s10856-012-4575-9

Nikravesh, N., Cox, S. C., Ellis, M. J., & Grover, L. M. (2017). Encapsulation and fluidization maintains the viability and glucose sensitivity of beta-cells. ACS Biomaterials Science & Engineering, 3(8), 1750–1757. doi:https://doi.org/10.1021/acsbiomaterials.7b00191

Pehlivanovic, B. (2020). Curcumin: natural antimicrobial and anti-inflammatory agent. J. Pharm. Res. Int, 32(43), 1–8.

Qi, M. (2014). Transplantation of encapsulated pancreatic islets as a treatment for patients with type 1 diabetes mellitus. Advances in medicine, 2014. doi:https://doi.org/10.1155/2014/429710

Ricci, M., Blasi, P., Giovagnoli, S., Rossi, C., Macchiarulo, G., Luca, G., Calafiore, R. (2005). Ketoprofen controlled release from composite microcapsules for cell encapsulation: effect on post-transplant acute inflammation. Journal of controlled release, 107(3), 395–407. doi:https://doi.org/10.1016/j.jconrel.2005.06.023

Roshanbinfar, K., & Salahshour Kordestani, S. (2013). Encapsulating beta islet cells in alginate, alginate-chitosan and alginate-chitosan-PEG microcapsules and investigation of insulin secretion. Journal of Biomaterials and Tissue Engineering, 3(2), 185–189. doi:http://dx.doi.org/10.1166/jbt.2013.1082

Samojlik, M. M., & Stabler, C. L. (2021). Designing biomaterials for the modulation of allogeneic and autoimmune responses to cellular implants in Type 1 Diabetes. (1878-7568 (Electronic)). doi:https://doi.org/10.1016/j.actbio.2021.05.039

Sarker, B., Papageorgiou, D. G., Silva, R., Zehnder, T., Gul-E-Noor, F., Bertmer, M., Boccaccini, A. R. (2014). Fabrication of alginate–gelatin crosslinked hydrogel microcapsules and evaluation of the microstructure and physico-chemical properties. Journal of Materials Chemistry B, 2(11), 1470–1482. doi:http://dx.doi.org/10.1039/C3TB21509A

Schaschkow, A., Sigrist, S., Mura, C., Barthes, J., Vrana, N. E., Czuba, E., Lejay, A. (2020). Glycaemic control in diabetic rats treated with islet transplantation using plasma combined with hydroxypropylmethyl cellulose hydrogel. Acta Biomaterialia, 102, 259–272. doi:https://doi.org/10.1016/j.actbio.2019.11.047

Skrzypek, K., Groot Nibbelink, M., Van Lente, J., Buitinga, M., Engelse, M. A., De Koning, E. J., Stamatialis, D. (2017). Pancreatic islet macroencapsulation using microwell porous membranes. Scientific reports, 7(1), 1–12. doi:https://doi.org/10.1038/s41598-017-09647-7

Sookkasem, A., Chatpun, S., Yuenyongsawad, S., & Wiwattanapatapee, R. (2015). Alginate beads for colon specific delivery of self-emulsifying curcumin. Journal of Drug Delivery Science and Technology, 29, 159–166. doi:https://doi.org/10.1016/j.jddst.2015.07.005

Toda, S., Fattah, A., Asawa, K., Nakamura, N., N Ekdahl, K., Nilsson, B., & Teramura, Y. (2019). Optimization of islet microencapsulation with thin polymer membranes for long-term stability. Micromachines, 10(11), 755. doi:https://doi.org/10.3390/mi10110755

